# Trajectory-Based Parameterization of a Coarse-Grained Forcefield for High-Thoughput Protein Simulation

**DOI:** 10.1101/169326

**Authors:** John M. Jumper, Karl F. Freed, Tobin R. Sosnick

**Affiliations:** University of Chicago

## Abstract

The traditional trade-off in biomolecular simulation between accuracy and computational efficiency is predicated on the assumption that detailed forcefields are typically well-parameterized (i.e. obtaining a significant fraction of possible accuracy). We re-examine this trade-off in the more realistic regime in which parameterization is a greater source of bias than the level of detail in the forcefield. To address parameterization of coarse-grained forcefields, we use the contrastive divergence technique from machine learning to train directly from simulation trajectories on 450 proteins. In our scheme, the computational efficiency of the model enables high accuracy through precise tuning of the Boltzmann ensemble over a large collection of proteins. This method is applied to our recently developed *Upside* model [1], where the free energy for side chains are rapidly calculated at every time-step, allowing for a smooth energy landscape without steric rattling of the side chains. After our contrastive divergence training, the model is able to fold proteins up to approximately 100 residues *de novo* on a single core in CPU core-days. Additionally, the improved *Upside* model is a strong starting point both for investigation of folding dynamics and as an inexpensive Bayesian prior for protein physics that can be integrated with additional experimental or bioinformatic data.

---

A major challenge in protein chemistry is to extract the underlying interaction energies from a set of proteins that capture the physiochemistry that lead to their folded structures. We address this challenge by showing that a strong connection exists between properties of the native basin and the rest of the protein's conformational landscape, and this connection is strong enough to train a potential for *de novo* folding simulations. Furthermore, the resulting potential is inexpensive enough to equilibrate simulations of small proteins in CPU core-days on a commodity computer.

Since Anfinsen’s original demonstration that a protein’s sequence determines its structure, multiple computational strategies have been developed to predict a protein’s structure from its sequence. An additional facet of this challenge is to replicate the energy landscape that defines both the folding process and other dynamical properties. In the absence of other information, coarsegrained models with one or a few beads per residue are too simplistic for *de novo* structure prediction. C_*β*_ level models having authentic protein backbones with *ϕ*/*ψ* dihedral angles, but lacking side chain rotamers, have achieved some success [2,3,4]. Within the last decade, allatom, explicit solvent methods have become successful for the folding of some small proteins, although the ability to replicate the properties outside the native basin requires improvement [5]. For the folding process, it is unclear which representation provides the optimal combination of detail and computational expense to replicate protein folding and dynamics. Integral to the choice of representation is the choice of interactions to include, such as hydrogen bonding, van der Waals interactions and hydrophobic burial.

Another factor to consider is the need for the training algorithm to balance the influence of all interactions. Protein thermodynamics reflects a delicate balance between the free energy of the folded and unfolded states. If one interaction term in the potential is slightly too large, the entire landscape can be severely distorted. For example, if backbone hydrogen bonding energies are too large compared to backbone-solvent interactions (which includes hydrogen bonds between the backbone and water), an excess of hydrogen bonding will ensue and pathways will be dominated by unrealistically stable native-and non-native secondary structures. In an extreme situation, long helices involving all residues could be the lowest energy structure.

The balancing of these various energies has been a major effort, and the balance is continually being adjusted as new forcefield biases are identified [6]. However, the adjustment of some parameters to correct one deficiency can inadvertently degrade performance for other quantities. In order to achieve the correct balance, all terms in the model should be trained together, rather than adjusted with an ad hoc procedure to correct each identified deficit.

To achieve this balance with a detailed interaction model, we use our recently developed, extremely rapid *Upside* implicit solvent molecular dynamics program [1]. In *Upside*, each residue is represented with a polypeptide backbone and a side chain interaction site or bead which can adopt up to 6 positions representing up to six different side chain *χ*_1_/*χ*_2_ states. The key advance of the model is the smoothing of the energy surface by approximate analytic integration of free energies for the side chains’ discrete states. When trained to predict side chain conformations from the Protein Data Bank (PDB), the method can fold a few small proteins with moderate accuracy in a core-day. The majority of speedup of the algorithm is a result of a unique side-chain calculation which directly calculates the side chain probability distribution and its free energy. This free energy calculation, performed at every time step, avoids the steric rattling of the side chains which can occur in the condensed phase in all-atom simulations, and so allows the backbone to move in a smoother energy landscape.

Here, we demonstrate that we can achieve *de novo* folding for a diverse collection of proteins by combining our fast-equilibrating *Upside* model with a contrastive divergence procedure that optimizes the accuracy of the native well. The resulting parameters are sufficiently balanced and accurate to achieve reversible folding for many proteins in our validation set. Furthermore, we demonstrate that gradient descent on energy terms using only data from sampled trajectories is sufficient to parameterize a protein model with tens of thousands of parameters. In addition, the resulting model is an excellent starting point for large scale protein simulations using more detailed models as well as the integration of large quantities of external information (such as contact predictions).

## COARSE-GRAINED MODEL

In our recently-developed *Upside* model, only the N, C_*α*_, and C atoms for each residue undergo dynamics. This simple representation of the protein allows for molecular dynamics on a smooth landscape but also makes it challenging to include the entirety of the protein physics. To address this challenge, we build additional layers of derived coordinates during the energy computation, much like virtual sites in a traditional force field. These layers include amide hydrogens, carbonyl oxygens, hydrogen bonding and residue burial scores, and the possible locations of protein side chains. All of the derivative information required is backpropagated through these layers of representation during the force computation for molecular dynamics. The most challenging representation is the side chain positions because a side chain packing problem must be solved in order to determine the distribution of their positions for a given backbone geometry. To pack the side chains probabilistically and obtain a side chain free energy, we use a self-consistent iteration as described in our recent work [1] (Fig. 1).

**Figure 1.**
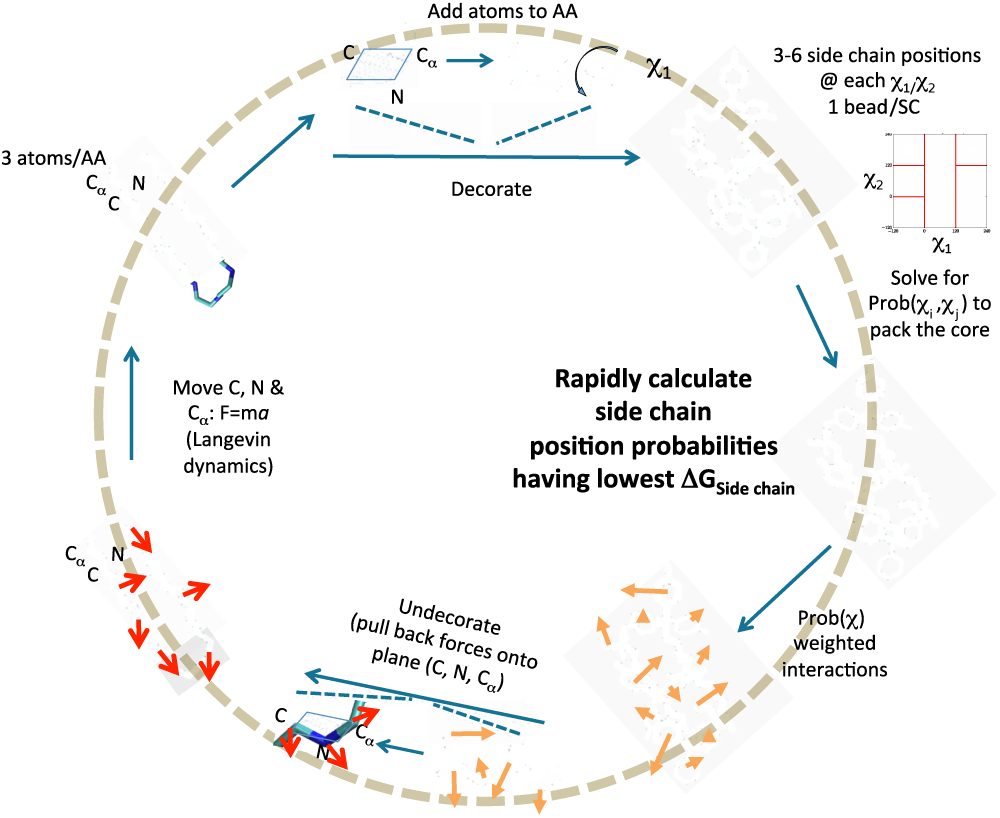
Computational inner loop for *Upside.* The possible positions of the protein side chains are added during each energy or force computation, then an approximate Boltzmann distribution is estimated for the side chains, and the free energy of the side chains is computed using the approximate Boltzmann ensemble. The resulting energy derivatives are pulled back to the backbone coordinates to update the backbone momenta.

The majority of *Upside* interaction parameters define the pairwise interactions between side chains, where each side chain is represented by a single directional bead. All of the pairwise interactions have the functional form

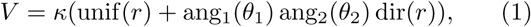

where unif, ang_1_, ang_2_, and dir are arbitrary curves represented by cubic splines. The potential for each of the 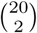 + 20 = 210 types of amino acid pairs are described with 62 spline coefficients per pair, giving 13020 parameters. There are also five interaction sites on the backbone, roughly representing the H, O, N, *C_α_*, and C atoms, with 54 parameters per interaction due to a smaller cutoff radius. The total number of side chain-backbone interaction parameters is 5400.

We add an additional term to capture desolvation effects by computing the number of side chains within a hemisphere above the C_*β*_ (a derived position from the backbone positions). To handle the uncertainty of rotameric states that can affect the count, the count for different rotameric states are weighted by the prior probabilities of the rotamer states, given by

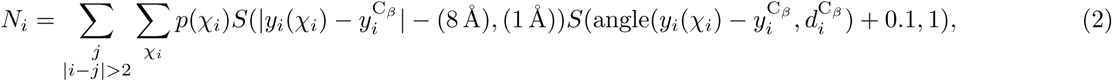

where *S* is sigmoid-like cutoff function, *y*^C_*β*_^ the position of the C_*β*_, and *d*^C_*β*_^ is the C_*α*_–C_*β*_ bond direction. High values of *N_i_* correspond to buried residues and low values correspond to exposed residues. The total energy is

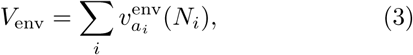

which is the sum of the individual 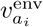 potential curves for each residue *i*. While more sophisticated solvation potentials exist, our implementation is very fast and easily optimized by the contrastive divergence procedure, while remaining flexible enough to represent many of the solvation effects omitted by the two-body side chain potential.

The backbone Ramachandran potential is 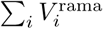(*ϕ_i_*,*ψ_i_*), where 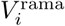 depends on the chemical identity of the *i* – 1, *i*, and *i* + 1 residues. The Ramachandran potentials are based on the turn, coil, or bridge (TCB) Ramachandran probability models in the NDRD backbone library [7]. We introduce a single parameter controlling extra stabilization of angles consistent with, *β*-sheet geometries to allow training to counteract an observed tendency for our model to overstabilize helices. The backbone non-bonded interactions are governed by a distance-and angle-dependent hydrogen-bonding potential whose magnitude (but not geometry) is chosen by contrastive divergence. The backbone N, C_*α*_, C_*β*_, and C feel a steric repulsive interaction at approximately 1.5 Å.

## CONTRASTIVE DIVERGENCE METHOD

Our implementation of contrastive divergence considers two ensembles, one closely restrained to the native (crystal) structure and another that is free to diffuse away during simulations. In a perfect model, an unrestrained ensemble would remain close to the native structure. For an inexact model, differences will arise, such as an excess of backbone-backbone hydrogen bonding in the free ensemble. Reducing the hydrogen bond energy would shift the free ensemble closer to the native ensemble. The parameter modification must be small, however, because shifting the hydrogen bond energy may adversely affect other parameters, e.g., by reducing the burial energy. Accordingly, after each set of simulations is run on a subset of our training set, we modify all the parameters with small updates to shift the simulation ensemble to better match the native-restrained ensemble. The algorithm is converged when no parameter can be altered to shift the free ensemble closer to the native-restrained ensemble.

The free ensemble is generated using 5000 time units of dynamics (approximately 10 wall-clock minutes), with the first half being discarded as equilibration. Unless the native state is particularly unstable, this time is insufficient for exploration of the conformational landscape much beyond the native basin (RMSD within 6 Å) and so produces only a locally-equilibrated ensemble.

The native ensemble is traditionally defined as a single conformation with precise 3D coordinates. This *δ*-function distribution is problematic for proteins because they are dynamical molecules whose solution ensemble may differ from the crystal structure for multiple reasons, most importantly errors and packing artifacts in crystallography. To reduce the impact of these issues, we replace the exact ensemble structures with the ensemble restrained to be near the crystal structure, within approximately 1 Å RMSD. This procedure is analogous to the restrained equilibration of crystal structures required to prepare systems for all-atom molecular dynamics. To account for changing parameters, we apply the restrained relaxation on every optimizer step.

After generation of the free and native-restrained ensembles, we change the energy parameters *α_i_*, where *i* is the optimizer step, in proportion to the amount that the change can differentiate the two ensembles. This procedure is a form of gradient descent to reduce the “distance” between the free and native-restrained ensembles,

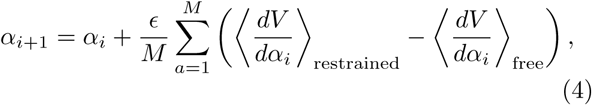

where *ϵ* the step size, *M* is the number of proteins, and *a* indexes the simulated proteins. The quantity 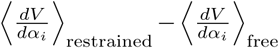 represents a pseudo-derivative of the free energy of restraining the simulation to be near the crystal structure. In the limit that the simulation duration is infinite, this difference is the exact derivative of the free energy. In practice, this difference chooses a suitable direction to improve the parameters.

The simulations use temperature replica exchange with eight replicas to enhance barrier crossing [8], while the temperature intervals of the replicas scale with 1/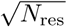 to encourage efficient replica exchange for proteins of various sizes. The progress of the replica exchange is monitored by the average RMSD-to-crystal structure over the simulation for each minibatch (subset of training proteins).

The initial parameters for the potential come from optimizing side chain packing accuracy of the model. The contrastive divergence training rapidly improves this model as there is a quick decline in average RMSD over a minibatch from 6 Å to 3 Å. This decline is accompanied by rapid movement of the parameters. To reduce parameter fluctuations and fine-tune the results, we reduce the optimizer step size by a factor of four after two full passes through the training set. While the slope of RMSD change with respect to the number of steps has greatly decreased over the iterations, there are indications that the parameters have not yet converged. Earlier tests, however, showed that continuing the contrastive divergence until convergence does not necessarily produce better results, as has been previously observed [9]. When large barriers surround the native states, minimal relaxation of the conformation occurs, which in turn provides little new information, and further finetuning may even *reduce* the accuracy of the model. Additionally, early termination of optimization has been observed to function as a regularizer that favors simpler models [10].

**Figure 2.**
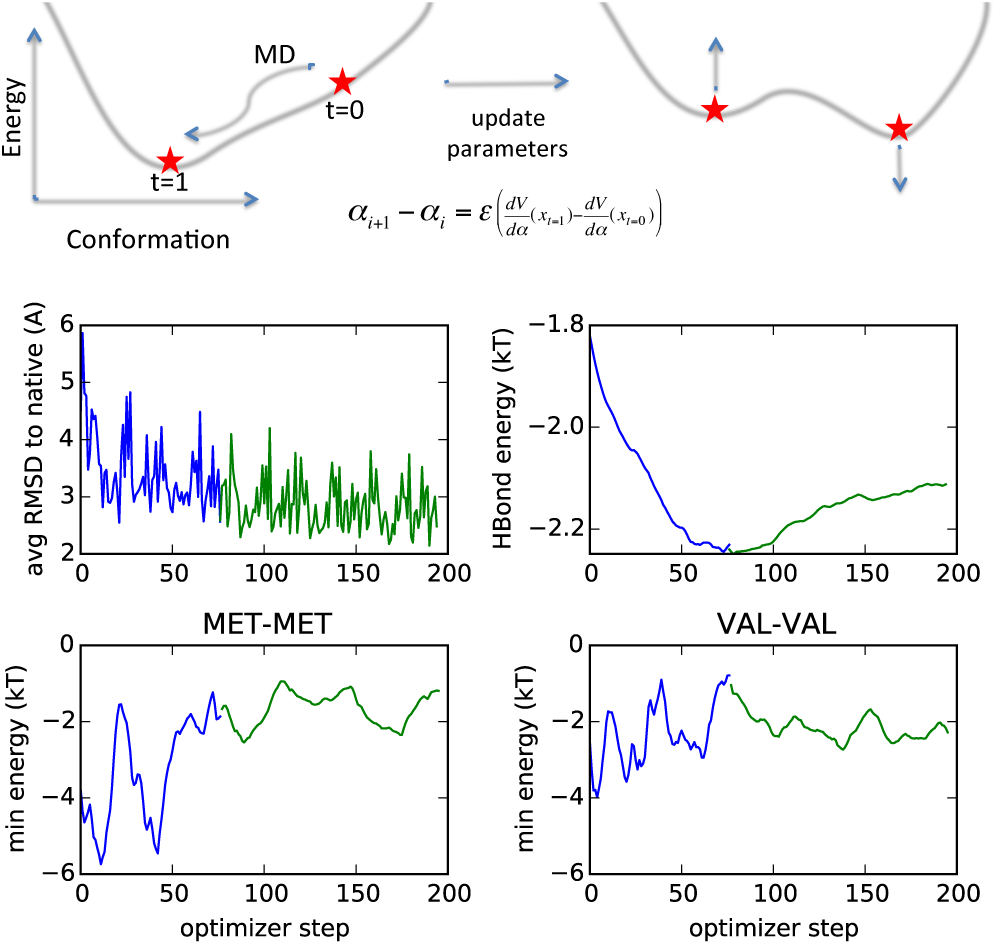
Cartoon and time-series of contrastive divergence training. In all plots, the blue curves indicate larger step-size training and the green plots indicate smaller step-size (finetuning). The upper left plot shows the decline in minibatch-averaged RMSD over the course of the optimization. The remaining plots show the convergence of the hydrogen bonding and side chain-side chain interaction parameters over the optimization. The larger step-size optimization of the side chain parameters exhibits large oscillations that inhibit convergence.

The hydrogen bond strength unexpectedly appears to converge to a significantly smaller value during the late, finetuning stage than during the early phase with larger optimizer steps. We speculate that the extra noise in the side chain interactions during the larger optimizer steps may in aggregate cause stronger side chain interactions for the protein. This effect would necessitate a large hydrogen bond energy to balance the increase in side chain interactions.

## ACCURACY OF STRUCTURE PREDICTION

Contrastive divergence training has been shown to train models well for many machine learning problems [11], even without having simulations that converge the Boltzmann ensemble. To test the accuracy of contrastive divergence on our protein model, we attempt *de novo* folding of a benchmark set of small, fast-folding proteins similar to those used in references [12, 13]. Before training, we remove homologous proteins from the training set to help ensure that this would be a true *de novo* prediction.

Two replica exchange simulations are run for each protein. The first set is initialized from the native configuration to assess the stability of the experimental structure. The second simulation is initialized from an unfolded state with Ramachandran *ϕ* and *ψ* angles chosen at random. The range of temperatures were chosen to be large enough to cover the unfolding transition.

We judge the accuracy and equilibration from the histogram of best-fit RMSD deviations from the native structure after discarding the initial third of the simulation as equilibration (see Fig. 3). When the native-initialized and unfolded-initialized structures have similar RMSD distributions, the simulations are likely converged. Proteins such as *α*3d and WW are approximately converged by this criterion but protein L and ubiquitin are not.

**Figure 3.**
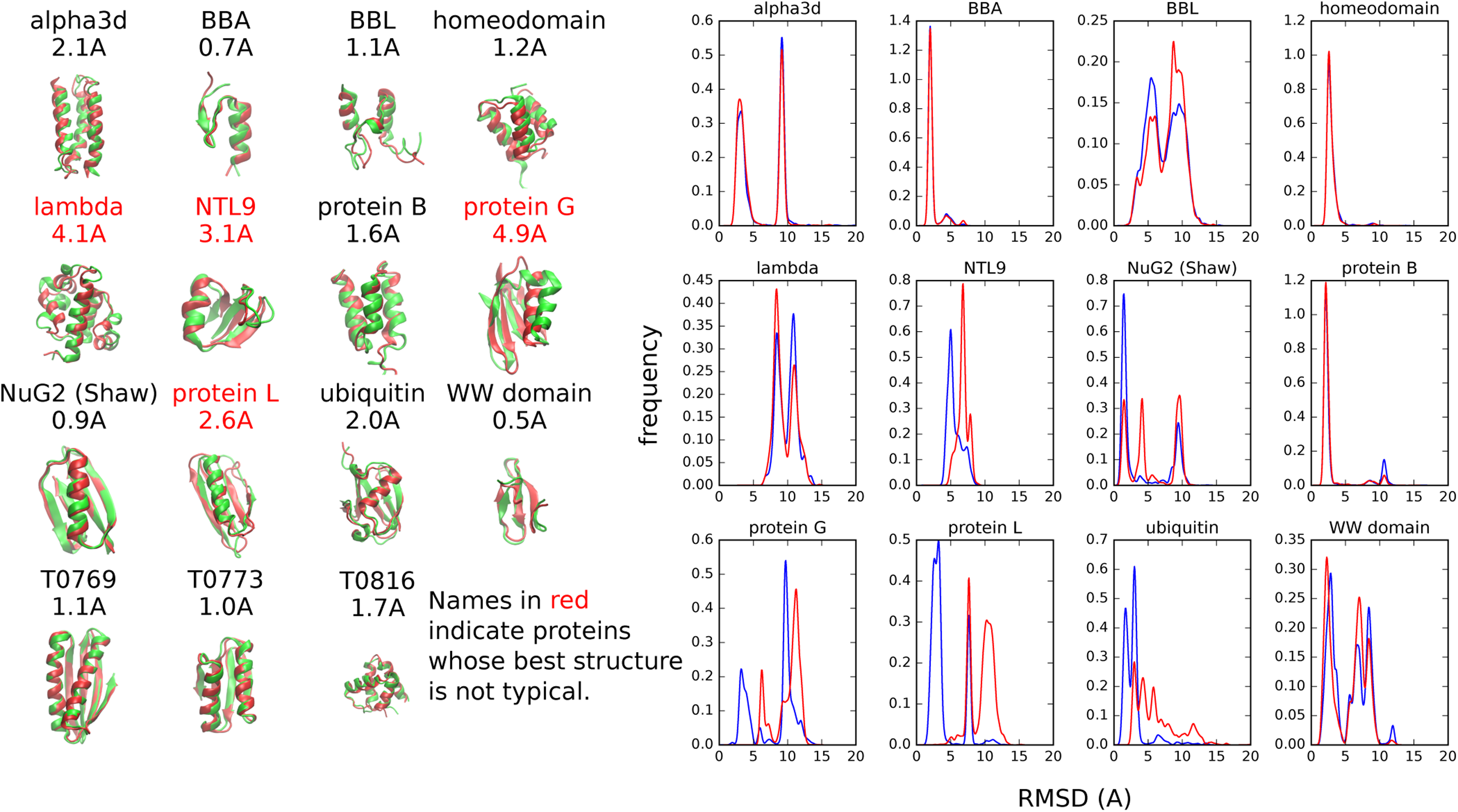
RMSD distributions and best structures after equilibration phase (see section for details). The simulations start from either the native (blue) or a random unfolded state (red). Simulations at each temperature were run for about three days with one CPU-core per replica.

The majority of the proteins show a small number of well-defined and stable basins that represent the dominant conformations with the current potential. While the simulations often produce several conformations quickly, equilibration of their populations takes longer (on the order of CPU-days for some proteins, though still extremely short in comparison to typical molecular dynamics simulations).

The *Upside* simulations tend to achieve the correct secondary structure with a small number of distinct tertiary arrangements. This diversity in tertiary structures occurs as mirrored three helix bundles for *α*3d and protein B, as well as the subtle re-arrangements of NuG2. As these structures coexist with similar probabilities at low temperature, we hypothesize that the short-time contrastive divergence we are using does not provide a sufficient library of large changes in the tertiary structure to enable the potential to properly distinguish the various, similar conformations. This issue will be addressed in future studies.

## CHARACTERIZATION OF FOLDING BEHAVIOR

In constant temperature simulations, we observe reversible folding to the native state for a number of proteins in our test set in core-days, Figs. 4 and 5. The time scales of folding indicated by these trajectories imply that the time scales we employed in the contrastive divergence simulations are far less (often a factor of 100 or more) than required to equilibrate these proteins, implying that contrastive divergence is optimizing only over fluctuations in or near the native well.

**Figure 4.**
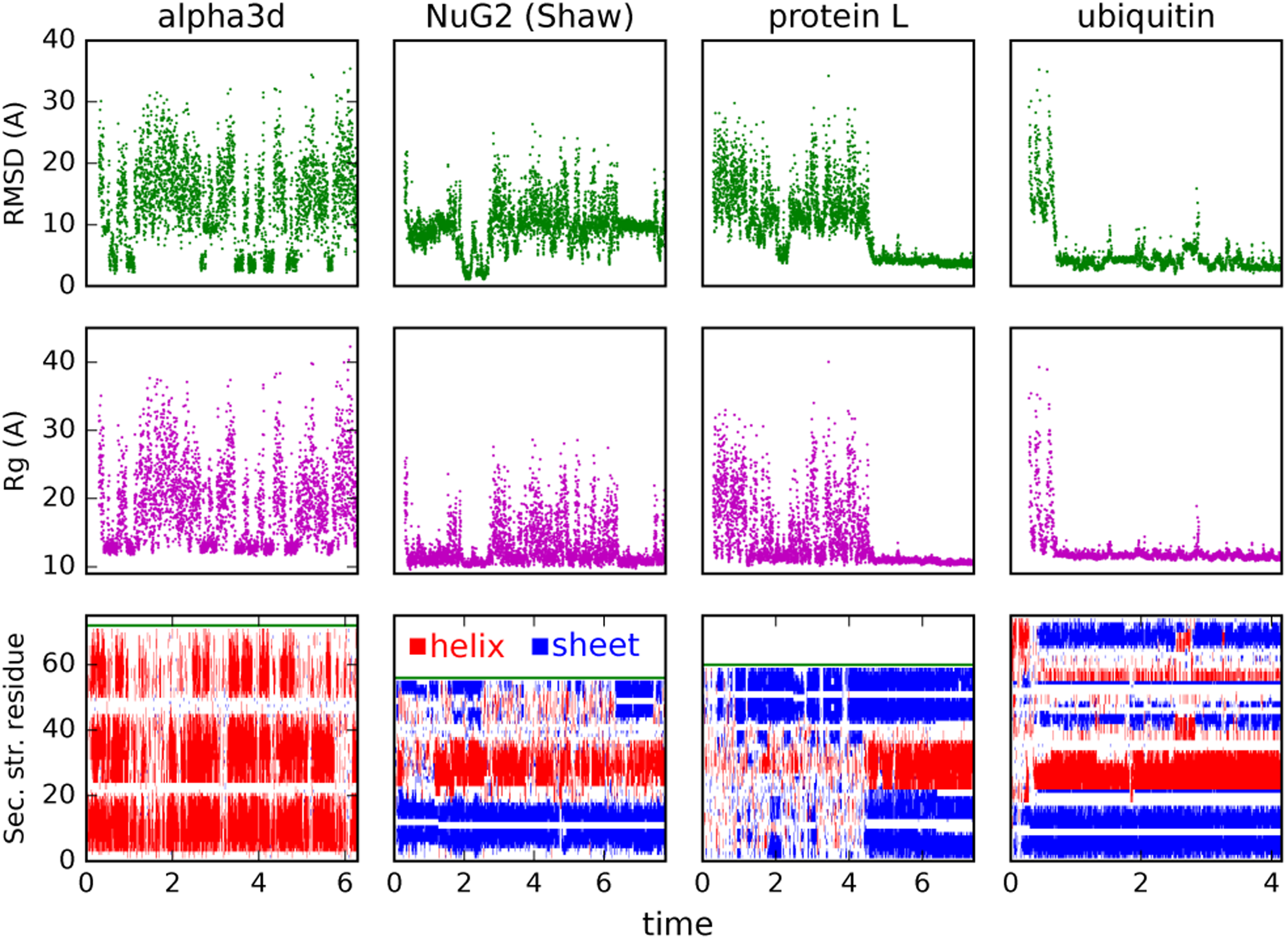
Constant temperature simulations. Trajectories are selected by the highest temperatures that still produce a significant population for the native state. Note that pivot Monte Carlo moves are attempted periodically which has little effect on folded dynamics but greatly decreases correlation time in the unfolded state.

**Figure 5.**
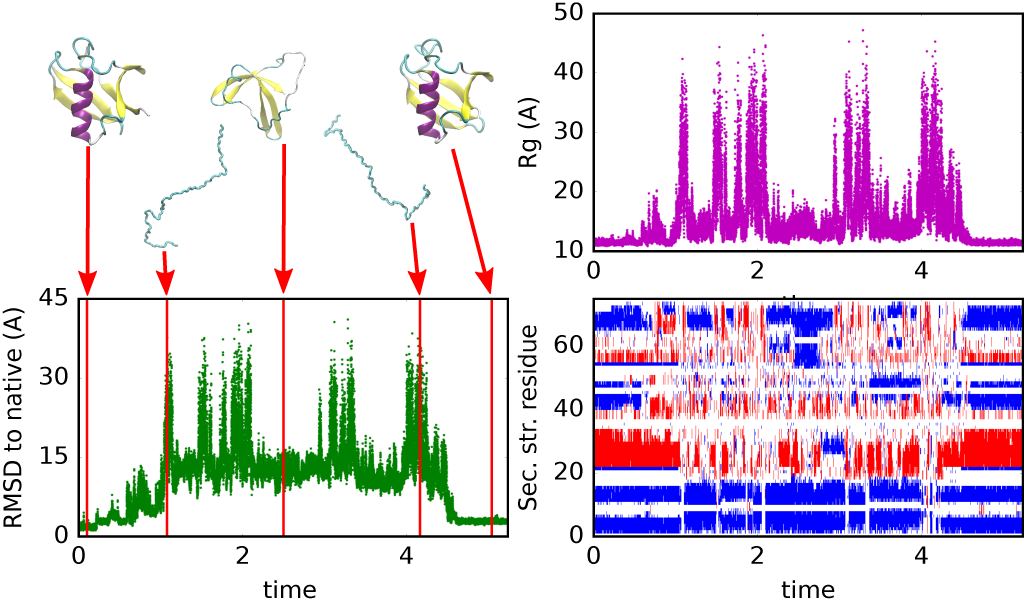
Constant temperature trajectory of ubiquitin at T=1.00, initialized from the native structure, with representative structures along the trajectory highlighted. The 2nd and 4th structures are chosen for having a high *R*_g_ while the last structure is chosen based on minimum RMSD (2.3 Å.) after achieving full unfolding.

Note that conditional on low hydrogen bonding, the radius of gyration (*R*_g_) at high temperature and at the peak of the heat capacity are quite similar. This suggests the increase in *R*_g_ for the unfolded state as temperature increases is driven by a reduction in backbone-backbone hydrogen bonds rather than side chain effects.

Based on these results, two observations should be reconciled. The first observation is the presence of a sharp phase transition with a single peak for the heat capacity. The shape of the phase transition, but not its amplitude, is consistent with a cooperative folding transition. The second observation is the unrealistically large level of residual hydrogen bonding in the denatured state at temperature of the maximum in the heat capacity. Although the hydrogen bonding is less than that in the native state, the residual hydrogen bonding indicates that the transition is not fully cooperative. These observations may be explained by the essential feature of the contrastive divergence process, that it must balance the competing energy terms of the model so that no one energy dominates. A small improvement to the contrastive divergence training may be able to push the temperature of melting secondary structure lower so that the folding is significantly more cooperative.

The *Upside* model exhibits concerted melting behavior over a small range of temperatures (see Fig. 6). While the temperature of the model in *Upside* is not exactly comparable to a physical temperature, it is reasonable to assume *T* =1 corresponds roughly to a temperature of 310 K. The ubiquitin transition occurs over a temperature range of approximately 0.07 temperature units, approximately a 20 K range, similar to that observed experimentally [15].

**Figure 6.**
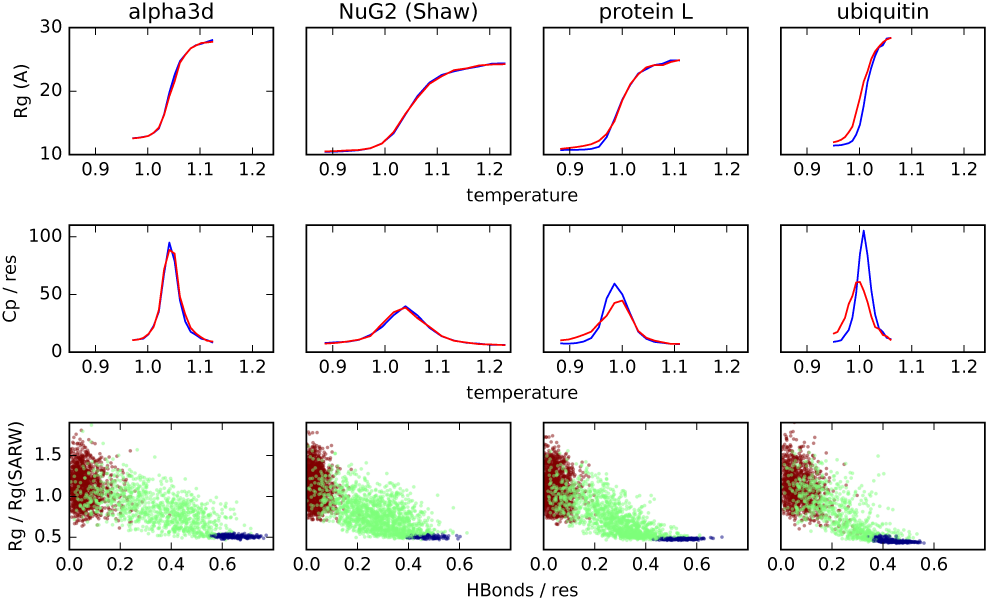
Thermodynamic behavior. The heat capacity is computed using the fluctuation relation *C*_p_ = (var *E)/T*^2^. In the bottom panels, the self-avoiding random walk *R*_g_ is computing using the experimental fit *R*_g_ = (1.9 Å)(*N*_res_)^0.6^ for chemically-denatured proteins [14]. The brown points are from high-temperature simulations, while the green and blue points are taken from simulations at the peak of the heat capacity for each protein — folded (blue) and unfolded states (green).

Furthermore, our temperature-denatured states have high *R*_g_ near the midpoint of the transition, consistent with experimental results and inconsistent with many allatom molecular dynamics folding simulations [5, 16]. At the peak of the heat capacity, the *R*_g_ is ~15% under the predicted from experimental data while the *R*_g_ at high temperature is ~10% above. Both *R*_g_ values are significantly larger than those in most atomistic molecular dynamics simulations [5].

## RELATED WORK

Contrastive divergence optimization has been applied to Gō-like protein potentials sampled with crankshaft Monte Carlo moves [17, 18]. These works optimized only tens of parameters, and the resulting model was used to fold protein G and 16-residue peptides.

Other studies have trained protein energy functions using libraries of decoys [19]. Such efforts are challenging because atomic energy functions have rugged energy landscapes where even small structural differences can produce large energy differences. This ruggedness implies that scoring decoys by energy without first relaxing them is problematic for the sharply-defined forcefields necessary to describe protein physics. This suggests that the best decoy set may be obtained instead by sampling trajectories of the protein energy function.

A distinction between contrastive divergence and traditional training methods, such as Z-score optimization [20], relates to the goal and the source of the decoys. In contrastive divergence, the critical task is to produce a high population of low RMSD structures with the model. Z-scoring training attempts to make the energy of the native state much lower than the average energy of of an pre-constructed decoy library. This is problematic because the decoys may not have structures that exhibit the pathologies of a poorly-trained model. Additionally, we believe optimization should concentrate on the lowest energies that have significant Boltzmann probability, not the average energy which is dominated by highly-unlikely structures. Furthermore, it is difficult to evaluate the reliable energies of decoys without relaxing the decoys. Methods based on simulation ensembles and the associated probability density (such as maximum likelihood and contrastive divergence) are well-defined and do not need pre-constructed decoy libraries.

Podtelezhnikov et al. [21] apply contrastive divergence to few-parameter protein models to optimize the parameters of hydrogen bond geometry. Their work is similar to this paper but narrower in scope.

Maximum likelihood requires the computation of the derivative of the free energy, which involves a summation over an equilibrium ensemble. Such a requirement necessitates a very long simulation to update parameters. Still, this approach can be viable when used with very small proteins on which the simulations converge quickly. A variant of maximum likelihood is given in [22], where decoys are generated and a maximum likelihood model is fit to adjust the parameters to distinguish between near-native and far-from-native conformations. The potential is trained on a single protein, tryptophan cage, and then the resulting potential is applied to a number of α-helical proteins with some success.

## DISCUSSION

We have developed a procedure involving extremely short simulations in the native energy well, coupled with optimization using contrastive divergence, to parameterize a sophisticated coarse-grain model. Underlying the model is a re-evaluation of the common assumption that increased detail is the path to greater accuracy. This requirement for detail is mitigated with trajectory-based training because less expensive models allow more extensive exploration leading to higher accuracy. We have also shown that very large numbers of parameters (even ∼20000 in our case) are no obstacle to producing accurate proteins models using trajectory-based training. While over-fitting is always a concern, the severity is greatly reduced because contrastive divergence is training *a*gainst the vast possibilities of alternative protein conformations explored by conformational sampling. Additionally, contrastive divergence automatically obtains balanced parameters such that no particular interaction overwhelms the others. We contend that this balance between parameters is more important than the accuracy of any particular term.

The precise time scale and temperature scale of the *Upside* models is intentionally left arbitrary because the coarse-graining process may leave us without a linear relationship to physical time and temperature. The speedup of *Upside* simulation due to the smoothing of side chain interactions is likely to have a disproportionate effect on time scales for collapsed structures as compared to extended structures. Regardless, the equilibrium population distribution that determines the free energy is expected to be approximately correct, as well as the order of dynamical folding events. The precise relationship of *Upside* time scales to physical time scales is left to future work.

Using a proper representation for protein physics is a key aspect of the *Upside* model. In particular, Upside decouples the representation of the protein used for dynamics, an N–C_*α*_-C backbone model, from the representation used for computing energies and forces, a sophisticated representation that includes oriented side chain interactions. This combination allows us to build up the sophisticated coordinates needed to represent solvent exposure of side chains, geometry of hydrophobic packing, and side chain-backbone hydrogen bonding without paying the cost of running dynamical simulation on a complex model with slow equilibrium in the condensed phase. The largest improvement is the application of belief propagation to the side chain degrees of freedom so that we represent detailed side chain physics at the χ_1_/χ_2_-level without incurring the roughening of the energy landscape and slowing of the dynamics normally associated with detailed sterics of side chain interactions. It is an open question to determine how much molecular detail must be retained for accurate protein energetics, but *Upside* gives us a flexible framework to explore these issues without compromising our simple backbone representation for dynamics.

## CONCLUSION

By employing a computationally fast yet detailed model, we can use multiple trajectories to train tens of thousands of parameters simultaneously to simulate protein folding and dynamics. The training successfully produces low-energy, native or near-native structures with sharp folding transitions for most of our validation proteins. The strategy’s success argues that simpler (in atomic representation) models that can be globally parameterized can rival more detailed but slower models whose parameterization is more challenging. Future work will address extending the timescale and size of the training. Coupling large computational resources with Markov state models [23] should improve training of the *Upside* model by exploring a larger and more diverse conformational landscape on each contrastive divergence step.

The ready generation of Boltzmann ensembles allows for a wide range of computational studies of protein folding, dynamics, and binding. For example, computational screening of large numbers of proteins for foldability should be tractable as is the study of hydrogen exchange and folding kinetics using computational alanine scanning. Additionally, in studies that incorporate experimental or bioinformatics data, including contact predictions, *Upside* provides an inexpensive Bayesian prior distribution over protein structures that may be updated using experimental information. This provides accurate predictions that make essential use of the totality of protein physics as encoded in the *Upside* model, while being inexpensive enough to allow validation and iteration on large numbers of proteins.

## MATERIALS AND METHODS

### Derivation of contrastive divergence

We derive the contrastive divergence method as a series of approximations to the problem of best approximating the probability distribution of observed PDB structures using a forcefield of an imperfect, fixed form. The initial part is a standard derivation of the maximum likelihood method, adapted to make clear its connection to protein molecular dynamics, while the end of the derivation makes clear the relaxation to obtain contrastive divergence as an approximation to maximum likelihood.

We begin by assuming that we have a large collection of protein sequences {*s_a_*} and their associated Boltzmann distributions 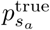(*X_a_*) under physiological conditions, where *a* represents an arbitrary label to enumerate the proteins and *X_a_* represents the configuration of the protein. Note that the “true” Boltzmann distribution is an unobservable idealization of the conformational ensemble of a protein under physiological conditions, and we further idealize that the true Boltzmann distribution is derived from from an extremely-complicated true potential 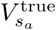 by statistical mechanics,

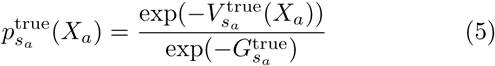

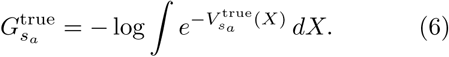

The subscript *s_a_* indicates that both the potential 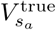 and free energy 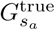 depend on the sequence of the protein. We may think of this as an artifact of working in the coarse-grained coordinates of the backbone trace, where the energy 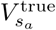 really represents the free energy of the backbone coordinates after integrating away the solvent and side chain degrees of freedom. An analogous situation occurs in parameterizing all-atom molecular dynamics, where the “energy” of the system really represents the free energy of the system after integrating over the electronic degrees of freedom. Our goal is to define a parametric 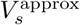(*X*) that approximates the 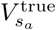 for any sequence *s*. We drop the subscript *s* below where there is no possibility for confusion.

For an approximate potential *V*^approx^, it is almost certain that *V*^approx^ does not have enough flexibility in its functional form to match all of the Boltzmann distributions *p_a_* for any sequence *s_a_.* We instead find a *V*^approx^ that is “close” to *V*^approx^. Defining the Boltzmann distribution of *V*^approx^ in the same manner as that of *V*^true^,

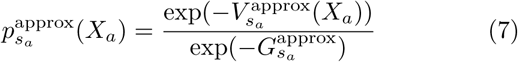

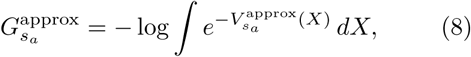

we may use the Kullback-Leibler (KL) divergence to measure the similarity of the associated Boltzmann distributions,

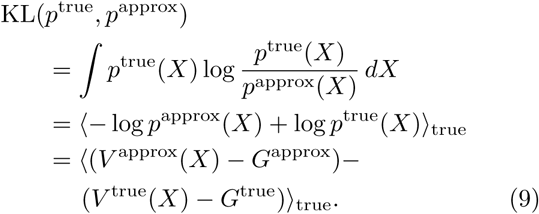

In the last equation, we note that the KL divergence is simply the average energy difference between the true and approximate potentials (after subtracting the free energies to normalize the probabilities), where the average is taken over the true Boltzmann distribution. The key fact when minimizing KL divergence is that if the approximate distribution lacks the freedom to exactly match the true distribution, then the minimizing distribution will be weaker than the true distribution (i.e. less sharp) to avoid assigning highly unfavorable energy to configurations that are likely in the true distribution.

Dropping constant terms, we may instead minimize

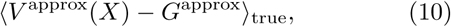

since the remaining term 〈*V*^true^(*X_a_*) – *G*^true^〉_true_ is independent of the approximating potential. This expectation value is still intractable since we do not know *p*^true^, but we can approximate,

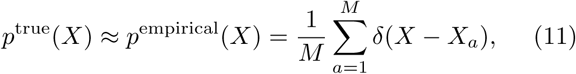

where *δ* is the Dirac delta function and *M* is the number of proteins. This gives the objection function,

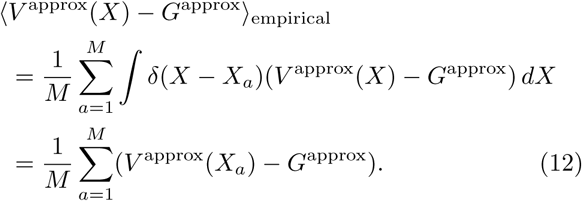

Minimize the expression [12] is exactly the method of maximum likelihood. The derivation given above illustrates two points via the connection to KL divergences. The first is that, if *V*^approx^ is insufficiently detailed, the model’s ensemble will be overly broad to ensure no experimental conformation has high energy under *V*^approx^. The second point is that with only a finite number of samples, *p*^empirical^ may be a poor approximation to *p*^true^, which would allow *V*^approx^ to wrap itself tightly near the *δ*-functions associated with each sample. This is the origin of overfitting in maximum-likelihood models.

We can now take the derivative with respect to an arbitrary forcefield parameter *α_i_* in preparation to perform gradient descent on [12]. The gradient is given by

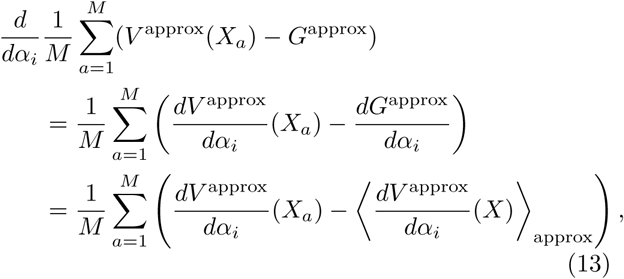

where we have used the standard statistical mechanics identity *dG*/*d_α_i__* = 〈*dV*/*d_α_i__*〉. While we have obtained a concrete expression for gradient descent in [13], we still have a major stumbling block. Computing the expectation of the derivative of the potential at *X_a_* is straightforward given a functional form for *V*^approx^, but obtaining even a reliable approximation for 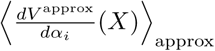 is extraordinarily difficult. To approximate the expectation value on a rough protein energy landscape, we would need Boltzmann samples from our current approximating potential. Even obtaining the single most likely configuration for our approximating potential is equivalent to finding the native state of the model, and this is very difficult for realistic pairwise potentials. Instead, we require the Boltzmann ensemble for all the proteins in our training set, and must update those Boltzmann ensembles as we use gradient descent to optimize the approximating potential. This represents an extreme expense and is unrealistic for anything but the simplest models of proteins. Note also that we cannot simply construct a large list of structures at some time and reweight those structures according to the potential, since the potential is constantly changing. Reweighting ensembles is only valid over very small neighborhoods of parameter space, and this procedure would depend on being able to generate an exhaustive survey of candidate structures in an exponentially large space.

The contrastive divergence method [24] approximates the maximum likelihood procedure using an empirical observation. We do not need an accurate approximation to [13], so long as the derivative points in direction of parameter space that improves the potential accuracy (i.e. any direction is acceptable as long as it is not uphill). Hinton proposes replacing Boltzmann average 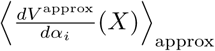 with a finite-time Fokker-Planck average over a very short period of time *for a simulation that originates at the data point X_a_*. In the Monte Carlo (MC) dynamics that the original authors use, even one MC step is sufficient to produce acceptable optimization of their model. In our case, we replace their small number of MC steps with a short time simulation using replica exchange Langevin dynamics. As the duration of the simulation is increased, our derivative estimate will converge to the true derivative [13]. Our paper empirically demonstrates that equilibrating each model within only a local region around the crystallographic native state is sufficient for a good folding model, so long as a large and diverse collection of protein structures are jointly optimized.

### Handling crystallographic artifacts

The derivation of contrastive divergence presented above makes the assumption that the conformations *X_a_* are equilibrium samples from the Boltzmann distribution of each protein, but in reality, we must work with crystal structures of proteins. While it has been shown that that the static diversity of crystal structures for different proteins conveys significant information about the dynamic ensembles of individual proteins [25]. Crystal structures deviate in a number of systematic ways from equilibrium samples, but we are most concerned about crystal packing artifacts, crystallizability bias, and errors in published structures. We expect that our bias in working only with crystallizable sequences, thus missing intrinsically disordered regions from training, likely biases the resulting potential to disfavor coil states. The loop-stabilizing effects of crystal packing somewhat counteract this effect, as it allows longer loop regions to exist in crystal structures.

We expect that our bias in working only with crystallizable sequences, thus missing intrinsically disordered regions from training, likely biases the resulting potential to disfavor coil states. The loop-stabilizing effects of crystal packing somewhat counteract this effect, as it allows longer loop regions to exist in crystal structures.

### Optimization and simulation details

The force is integrated using Verlet integration with a time step of 0.009 time units. Temperature is maintained using a Langevin thermostat with a thermalization timescale of 0.135 time units.

Simulation times in all figures are given in millions of *Upside* time units (approximately 10^8^ force evaluations).

The following temperature ranges are used for the replica exchange simulations in Figure 3 with 16 replicas per simulation. These temperatures are chosen to use the minimal temperature range that approximately span the thermal melting transition for each protein using information from an earlier set of replica exchange simulations. The temperatures of the simulation initialized from the crystal structure and those initialized from extended structures use the same temperature range.

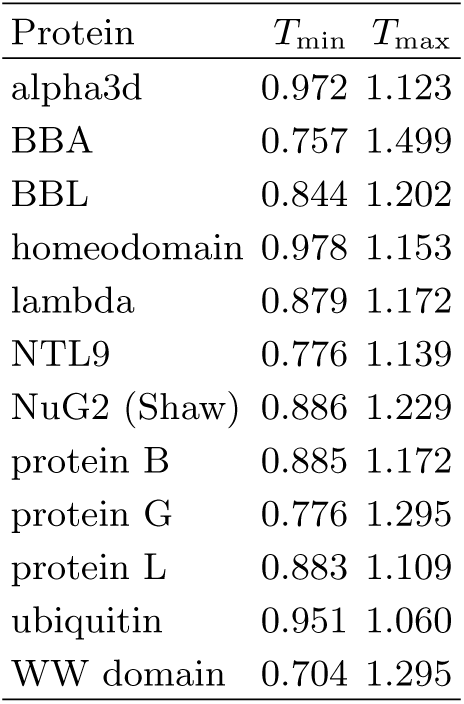

### Training data and optimization

The contrastive divergence training is conducted with 456 crystal structures from the Protein Data Bank. The initial selection of structures uses the PISCES server [26] to select proteins with X-ray resolution less than 2.2 Å and pairwise sequence similarity less than 30%. In structures with multiple chains, a single chain is chosen by the PISCES server. To avoid non-globular proteins or proteins with strong interactions with other subunits in the structure, random sample consensus linear regression [27] is used to identify outliers based on the relationship between log *N_res_* and log *R_g_*. Only chains with between 50 and 100 residues are used to encourage fast relaxation during the contrastive divergence simulations. All proteins homologous to proteins in the benchmark folding set are eliminated from the training set. Additionally, all proteins with backbone gaps, either missing residues due to diffuse electron density or non-standard amino acids that *Upside* does not handle, are also excluded from the training set.

The final training set of 456 proteins is divided into 38 groups of 12 proteins each, called minibatches. The Adam optimizer [28] is used to perform gradient descent on the objective function, using the contrastive divergence pseudo-gradient in place of the true maximum likelihood gradient. The Adam parameters used are *β*_1_ = 0.8, *β*_2_ = 0.96 and *ϵ* = 10^‒6^. The *α* parameter is varied based on the type of term to ensure stability, *α*_SC_ = 0.5, *α*_env_ = 0.1, *α*_HBond_ = 0.02, and *α*_heet_ = 0.03. The *α* parameters are multiplied by 0.25 for the finetuning optimization.

Regularization and derivative propagation for contrastive divergence optimization are handled using the Theano library [29].

### Details of test proteins

Mutations from the listed PDB structures are indicated in **bold**. The NuG2 sequence is from reference [12].

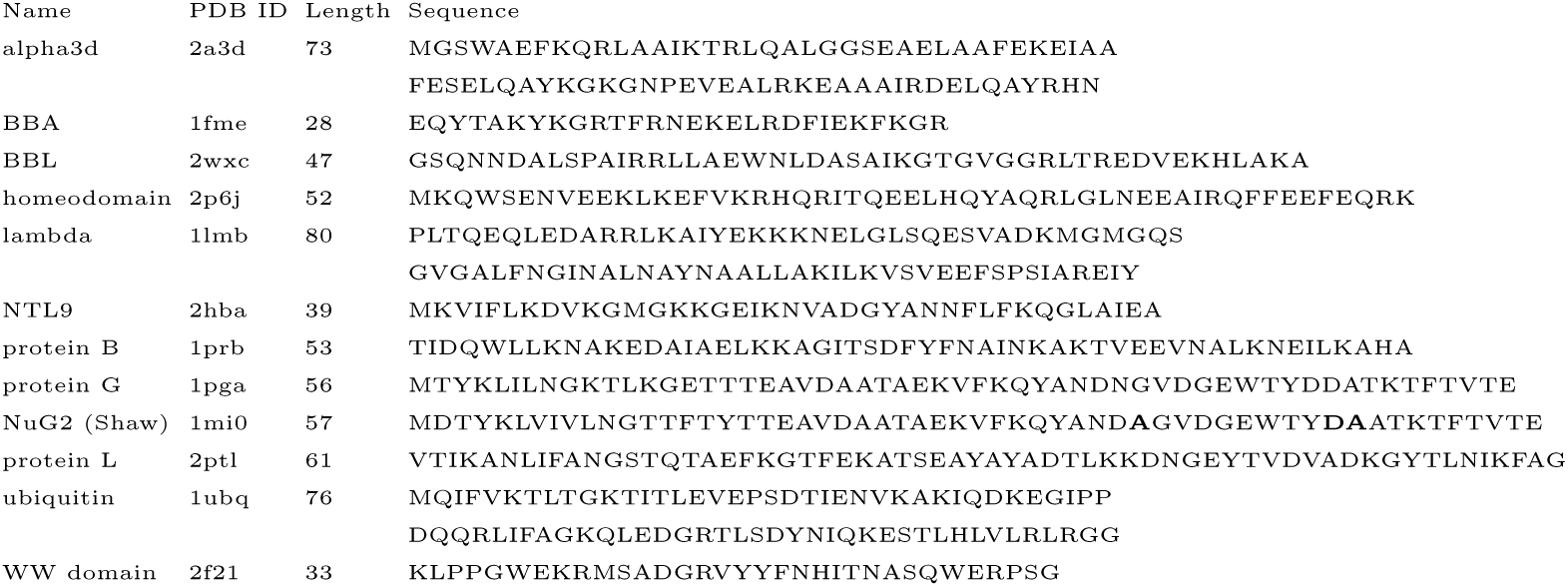

## ACKNOWLEDGMENTS

This research is supported, in part, by National Science Foundation (NSF) Grant No. CHE-1363012, NIH grant GM 55694, and by the NIGMS of the National Institutes of Health under award number T32GM008720. This work was completed in part with resources provided by the University of Chicago Research Computing Center. We would like to thank Sheng Wang and Jinbo Xu for helpful discussions during this research.

